# A triple functional sensing chip for rapid detection of pathogenic *Listeria monocytogenes*

**DOI:** 10.1101/2020.10.22.348615

**Authors:** Yachao Zhang, Huimin Wang, Sa Xiao, Xia Wang, Ping Xu

## Abstract

Here a triple functional sensing chip was created for *L. monocytogenes* detection by integrating three biomarkers (Listeriolysin O (LLO) at protein level, *hly* gene at genetic level, and acetoin at metabolic level). Liposome encapsulated catechol was used for LLO detection via LLO pore-forming ability. *hly* gene was specifically captured by using a thiolated capture probe on nanoporous gold (NPG). As an electroactive label, methylene blue was embedded in double-stranded structures to generate an electrochemical signal for *hly* detection. Combined with the electrocatalysis of NADH by NPG, the acetoin detection was achieved by measuring the consumption of NADH as a cofactor under acetoin reductase catalysis. Importantly, the *L. monocytogenes* detection results obtained by detecting three biomarkers using the chip can be mutually verified, which reduces the probability of false positives based on a single marker. Moreover, the detection time was reduced to about 90 min, making it a rapid and reliable tool for *L. monocytogenes* detection.

## Introduction

*Listeria monocytogenes*, a gram-positive bacterium, was first described in a disease that affected rabbits and guinea pigs in 1926^1^. It was recognized as the pathogenic cause of a human disease in the 1970s and later identified as a food borne pathogen in the 1980s^2^. Listeriosis is a kind of disease caused by the ingestion of foods that contaminated with *L. monocytogenes*, which mainly affects immunocompromised patients, pregnant women, and newborns^3^. After ingestion of contaminated foods, *L. monocytogenes* can cross intestinal barrier, blood-brain barrier as well as fetoplacental barrier, and cause gastroenteritis, meningitis, septicemia, abortion, and complications in pregnancy^3,4^. Therefore, the reliable and rapid detection of *L. monocytogenes* in foods is of considerable importance for preventing infections.

Extensive research in past decades has provided several detection methods for *L. monocytogenes*. Conventional culture methods based on colony morphology, sugar fermentation, and hemolytic properties are currently gold standards for the identification of *L. monocytogenes*^5^. However, these methods usually require 3~5 days to identify *L. monocytogenes*^6^, which does not meet the rapid detection requirements of food safety. Consequently, more rapid procedures utilizing enzyme-linked immunosorbent assay (ELISA)^7^, flow cytometry^8^, and molecular-based techniques (DNA hybridization^9^, and real-time quantitative polymerase chain reaction (qPCR)^10^) have been developed. Although these methods are quicker, they also have several drawbacks. For ELISA, the chromogenic procedure is complicated, and the chromogenic reaction takes a long time (at least 15min)^11^. Additionally, the use of specialized equipment, such as a microplate reader, limits the widespread application of ELISA-based detection in low-cost food industries. Although the sensitivity of molecular-based techniques is higher than ELISA, technical expertise as well as expensive chemicals and equipment are required for these techniques, which is the major limitation for their wider application^5^. Additionally, several rapid detection methods such as PCR and ELISA register false positives^12,13^. Thus, to ensure accurate detection, several different detection methods are simultaneously used in the identification of *L. monocytogenes*. However, the expertise and equipment required for different detection technologies vary, which substantially increases the difficulty and cost of *L. monocytogenes* detection. In view of the importance of *L. monocytogenes* detection and the disadvantages of these available methods, there is an urgent need to build a multi-biocatalyst analysis system that can integrate different methods for *L. monocytogenes* detection.

Biosensors have attracted wide attention because of their unique advantages such as continuous on-line detection, low cost, high selectivity, and fast response^14^. Further, different substances can be detected using a single biosensor^15,16^. Therefore, a multi-biocatalyst analysis biosensor that integrates different methods may be a good choice for *L. monocytogenes* detection. Among traditional molecular-based detection methods, biosensors based on nucleic acid markers are an alternative method. Because target DNA can be caught by a capture probe immobilized on an electrode via base-pairing interactions^17^, good specificity and a fast response are ensured. Then, an electroactive label such as methylene blue (MB) embedded in the double-stranded structure can serve as an electroactive substance to generate an electrochemical signal^18^. To detect *L. monocytogenes* using a nucleic acid marker-based biosensor, the selection of a specific nucleic acid marker is crucial. The specific nucleic acid marker should be a unique DNA or RNA carried by each species of the pathogenic microorganism that can be used to differentiate it from other organisms. *hly* gene, a virulence gene encoding Listeriolysin O (LLO), has been commonly used as a marker gene for *L. monocytogenes* qPCR detection^19,20^. Thus, a biosensor based on the *hly* gene can facilitate the rapid detection of *L. monocytogenes*. Besides, LLO protein is a pore-forming toxin^21^. It can bind to the cholesterol of host membranes, oligomerize, and form large pores to help *L. monocytogenes* escaping from the host vacuole^22^. Therefore, the pore-forming ability of the LLO protein can be used to build a liposome biosensor for the detection of the LLO protein. Besides, it has been reported that acetoin, known as 3-hydroxy-2-butanone, could serve as a marker for the detection of *L. monocytogenes*^23^. The concentration of acetoin showed a good positive correlation with the quantity of *L. monocytogenes*^23,24,25^. Based on previous research, acetoin can be converted to 2,3-butanediol by acetoin reductase (AR) in the presence of NADH as a cofactor^26^. Thus, the detection of acetoin can be achieved by measuring the consumption of NADH. The integration of these different markers may provide the possibility for rapid and accurate detection of *L. monocytogenes*. However, a single sensing chip integrated multiple markers for pathogenic *L. monocytogenes* detection remains an unmet challenge due to the different requirements of different markers.

In the current study, a triple functional sensing chip was created for *L. monocytogenes* detection by combining a toxin marker (LLO), a nucleic acid marker (*hly* gene), and a metabolite marker (acetoin) on a multi-biocatalyst sensing chip. The liposome encapsulated redox substance was used for the LLO protein detection. In addition, a thiolated capture probe synthesized based on *hly* gene sequence was used to recognize *hly* gene, and MB was used as an electroactive label to generate an electrochemical signal. For acetoin detection, *bdha* gene encoding AR was overexpressed and purified. Nanoporous gold (NPG) was selected as an ideal immobilization material for the multi-biocatalyst sensing chip. Then, a triple functional sensing chip was constructed for *L. monocytogenes* determination by integrating a toxin marker (LLO), a nucleic acid marker (*hly* gene), and a metabolite marker (acetoin). The electrochemical performance of the triple functional sensing chip was evaluated in detail.

## Results

### Construction principle of the triple functional sensing chip

As for *L. monocytogenes* detection, traditional detection technology (such as PCR and ELISA) based on a single marker detection method may result in false positives^12,13^. To improve the accuracy of detection results, several different detection methods based on different markers are simultaneously used^27^. However, many issues are created by the simultaneous use of multiple detection methods, such as the long sample-to-answer time and limited prospects for lab-free use^28^. Therefore, it is necessary to develop a single sensing chip based on multiple-level markers (such as toxin marker at the protein level, DNA markers at the genetic level, and small molecules at the metabolic level) to achieve simple, rapid, and accurate detection of *L. monocytogenes*. Because an integrated sensing chip needs a variety of electrocatalysts and detection systems, each with its own fundamentally distinct sensing format and fabrication requirements, the construction of an integrated sensing chip is very complicated. The supporting material of the integrated sensing chip is the key to solve the above-mentioned technical difficulties. The ideal supporting material should have the following properties: 1) a large specific surface area which can provide a large amount of site for bio-recognition elements binding; 2) special electrocatalytic ability which can generate an electric signal by electrocatalysis a substrate; 3) high electrical conductivity which can accelerate electron transfer and guarantee electric signal strength. The screening of ideal supporting material may reduce the complexity of the integrated sensing chip construction.

In our previous work, it was found that NPG with bicontinuous three-dimensional nanoporous structure (as shown in Figure 1a and Figure 2a) was an ideal supporting material for the immobilization of bio-recognition elements such as enzymes or microbial cells^29,30,31^. Moreover, as a nanoporous metal material, NPG has an excellent electrical conductivity and efficient electrocatalytic activity towards many substrates, such as catechol^31^ and NADH^32^. These excellent properties of NPG can be utilized in the construction of the multi-biocatalyst sensing chip for pathogenic microorganism *L. monocytogenes* determination by combining three markers (LLO protein, *hly* gene, and acetoin). The preparation process of the multi-biocatalyst sensing chip was detailly described in Figure 1a. A paper sensor with four carbon working electrodes (CWE), one carbon counter electrode (CCE), and one reference electrode (Ag/AgCl) was used as the multi-biocatalyst sensing chip (Figure 1a). CWEs were firstly modified by NPG films, and then the CWEs modified with NPG served as working electrodes to immobilize different bio-recognitions.

**Figure 1.**
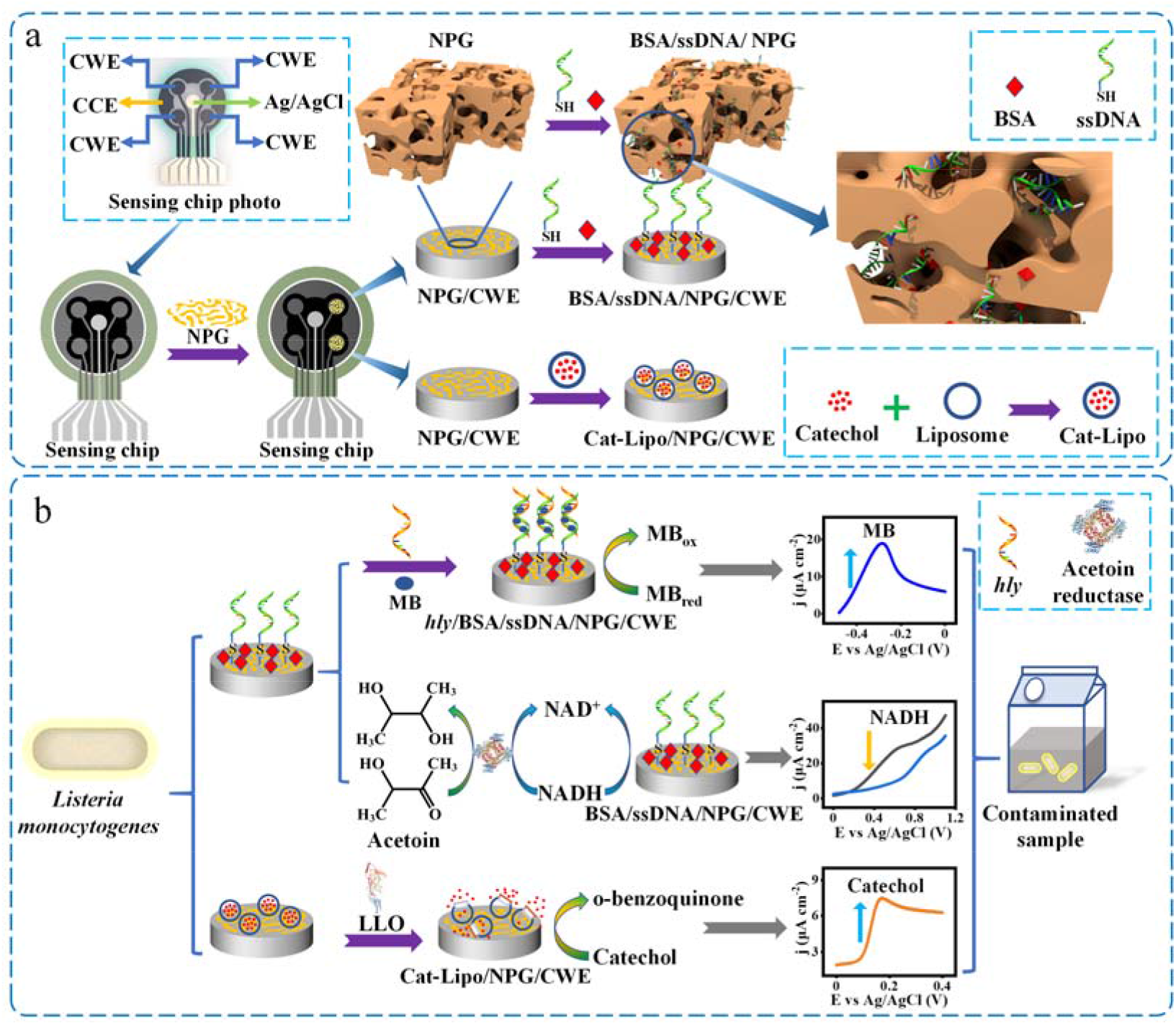
Construction principle of the triple functional sensing chip. (a) The three-dimensional diagram of NPG, and the preparation of the triple functional sensing chip (CWE: carbon working electrode; CCE: carbon count electrode). (b) The reaction mechanism of the triple functional sensing chip.

**Figure 2.**
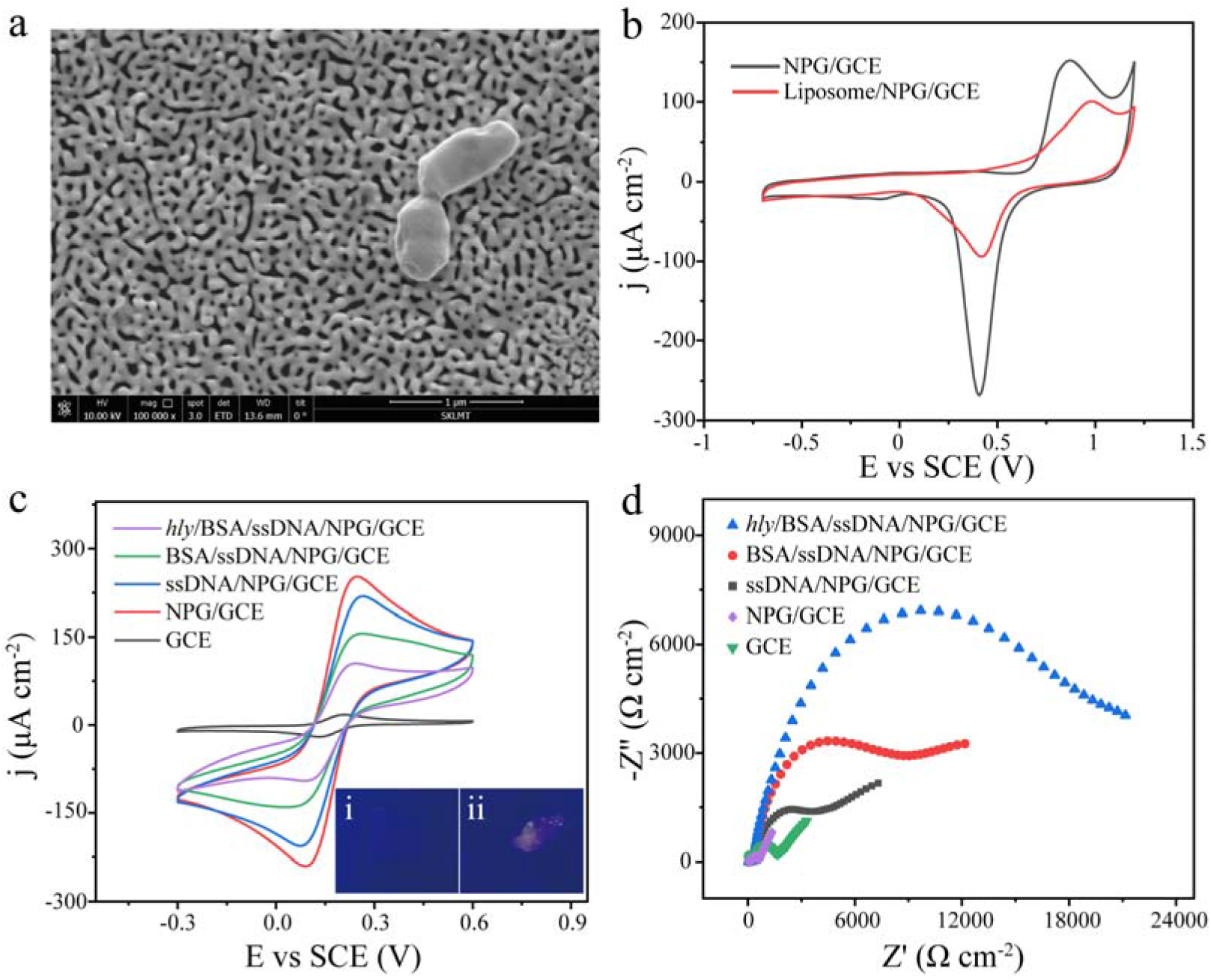
Characterization of the triple functional sensing chip. (a) The SEM image of the Cat-Lipo/NPG. (b) The CVs of the NPG/GCE electrode, and Cat-Lipo/NPG/GCE bioelectrode in a PBS (50 mM, pH 7.0). (c) The CVs of the GCE electrode, NPG/GCE electrode, ssDNA/NPG/GCE bioelectrode, BSA/ssDNA/NPG/GCE bioelectrode, and *hly*/BSA/ssDNA/NPG/GCE bioelectrode in PBS (50 mM, pH 7.0) containing 5 mM K_3_Fe(CN)_6_/K_4_Fe(CN)_6_; inset: the morphologies of NPG (i) and *hly*/ssDNA/NPG (ii) (both incubated with nucleic acid dye) under 470 nm blue light irradiation. (d) The EIS of the GCE electrode, NPG/GCE electrode, ssDNA/NPG/GCE bioelectrode, BSA/ssDNA/NPG/GCE bioelectrode, and *hly*/BSA/ssDNA/NPG/GCE bioelectrode in 5 mM K_3_Fe(CN)_6_/K_4_Fe(CN)_6_ solution at the potential of +0.13 V with a frequency range from 0.01 to 10^6^ Hz.

For LLO protein detection, catechol-liposome (Cat-Lipo) immobilized onto the NPG-modified CWE to form Cat-Lipo/NPG/CWE bioelectrode (Figure 1a). The pore-forming phenomenon will occur when LLO protein adsorbs on the membrane of the Cat-Lipo, which can cause catechol release from the Cat-Lipo (Figure 1b). Then, the released catechol can be catalyzed by NPG generating a current signal as shown in Figure 1b, and the working principle could be expressed as follows^31^.

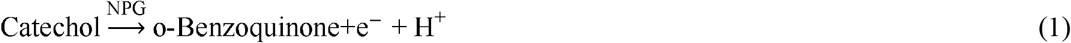

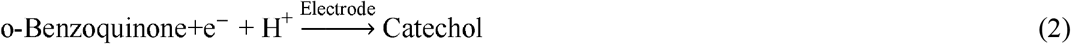

For *hly* gene detection, a thiolated capture probe (ssDNA), synthesized based on the *hly* gene sequence, was immobilized onto an NPG/CWE electrode via Au-S bond^29^, and bovine serum albumin (BSA) was used as a blocking reagent as shown in Figure 1a. After the preparation of the BSA/ssDNA/NPG/CWE bioelectrode, the target gene *hly*, which was complementary to the capture probe, bound to the capture probe immobilized on NPG, as shown in Figure 1b. As an electroactive label, MB intercalated into two successive base pairs of double-stranded structure^33^, and then generated an electrochemical signal (Figure 1b).

Additionally, it was reported that acetoin previously served as a marker for the detection of *L. monocytogenes*^23^. A positive correlation between of acetoin concentration and *L. monocytogenes* concentration was obtained^23,24,25^. Therefore, acetoin was selected as another marker for the detection of *L. monocytogenes* in this study. For acetoin detection, AR could catalyze acetoin reduce to 2,3-butanediol in the presence of NADH as a cofactor^26^. Combined with the electrochemical catalysis of NADH by NPG as mentioned above, the quantitative detection of acetoin was achieved by measuring the consumption of NADH using the BSA/ssDNA/NPG/CWE bioelectrode. Thus, the concentration of *L. monocytogenes* was calculated indirectly by the quantitative relationship between the production of acetoin and the concentration of *L. monocytogenes*. The working principle of the proposed the BSA/ssDNA/NPG/CWE bioelectrode for acetoin detection can be described as follows^34^.

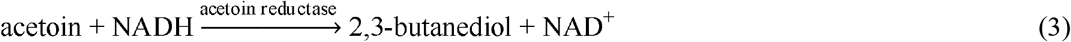

Based on these detection principles, the triple functional sensing chip was successfully constructed for pathogenic microorganism *L. monocytogenes* determination by integrating multiple detection markers onto a multi-biocatalyst sensing chip as shown in Figure 1.

### Characterization of bioelectrodes

In this study, different markers were detected using different bioelectrode on the integrated sensing chip. Therefore, these different bioelectrodes were characterized using different methods. To facilitate the study of electrode interface characteristics, bioelectrode CV characteristics, EIS analysis, MB signal generating by MB embedded in double-stranded structure, and synthetic *hly* detection were performed using modified NPG/GCE electrodes in a three-electrode cell as described in Experimental measurements.

First, the morphology of Cat-Lipo/NPG was characterized by scanning electron microscopy (SEM). As shown in Figure 2a, Cat-Lipo was successfully immobilized onto the surface of NPG. In addition, SEM revealed an open three-dimensional nanoporous structure of NPG with a pore size of ca. 35 nm, which could provide a higher surface area for the loading of capture probes and the free access of small molecule substrates such as catechol and NADH. Besides, Fujita et al. revealed that most gold ligaments of NPG had a hyperboloid-like shape with a negative curvature-accommodating nanopores and a positive curvature forming columnar ligaments^35^. These negative curvature-accommodating nanopores contained a high density of gold atomic steps which consisted of a large number of low-coordination atoms^36^. Low-coordination atoms can interact more strongly with molecules owing to their modified local electron structure, leading to reduced reaction barriers relative to gold surface^37^. Thus, this high density low-coordination atom can serve as an active site for the catalytic reaction, endowing NPG with active electrocatalytic ability.

To further confirm the immobilization of the Cat-Lipo on NPG, the cyclic voltammetry (CV) of the NPG/GCE electrode and Cat-Lipo/NPG/GCE bioelectrode were investigated in a phosphate buffer solution (PBS, 50 mM, pH 7.0) at a scan rate of 50 mV s^−1^. As shown in Figure 2b, the NPG/GCE electrode exhibited the highest redox peak current densities of NPG. In contrast, the redox peak current densities of NPG decreased after the loading of the Cat-Lipo. These results also confirmed that the Cat-Lipo/NPG/GCE bioelectrode was successfully constructed for the detection of LLO protein.

To confirm the immobilization of the capture probe on NPG, GeneGreen, as a nucleic acid fluorescence dye, was used as an indicator, which would produce fluorescence upon binding to double-stranded nucleic acids. After incubation with the nucleic acid fluorescence dye, the blue light irradiation morphologies of NPG and the *hly*/ssDNA/NPG composite were observed under 470 nm as shown in Figure 2c-i and Figure 2c-ii, respectively. It is important to note that only the nucleic acid fluorescence dye embedded into the nucleic acid chain can produce fluorescence under 470 nm blue light irradiation. Figure 2c-i shows that there was no fluorescence observed for NPG. Instead, the *hly*/ssDNA/NPG composite bound by GeneGreen emitted obvious fluorescence (Figure 2c-ii), which provided a morphological evidence for the successful immobilization of the capture probe.

Additionally, the GCE electrode, NPG/GCE electrode, ssDNA/NPG/GCE bioelectrode, BSA/ssDNA/NPG/GCE bioelectrode, and *hly*/BSA/ssDNA/NPG/GCE bioelectrode were characterized using CV in a PBS (50 mM, pH 7.0) containing 5 mM K_3_Fe(CN)_6_/K_4_Fe(CN)_6_ at a scan rate of 50 mV s^−1^. As shown in Figure 2c, a couple of distinct and symmetrical redox peaks of K_3_Fe(CN)_6_ were observed for all electrodes and bioelectrodes. After loading NPG onto the GCE electrode, the redox peak of K_3_Fe(CN)_6_ for the NPG-based electrodes or bioelectrodes were explored. The redox peak of K_3_Fe(CN)_6_ for the NPG-based electrodes or bioelectrodes were all found to be higher than that of the bare GCE electrode due to accelerated electron transfer by NPG^32^. Compared with the NPG/GCE electrode, decreases of peak current density were observed for the ssDNA/NPG/GCE bioelectrode, BSA/ssDNA/NPG/GCE bioelectrode, and *hly*/BSA/ssDNA/NPG/GCE bioelectrode. Moreover, the decreases of peak current density gradually increased with the loading of ssDNA, BSA, and *hly* gene (Figure 2c). The reason was that interfacial electron transfer was obstructed by the loading of ssDNA, BSA, and target gene *hly* on NPG due to their insulating nature. Accordingly, the redox peak currents of K_3_Fe(CN)_6_ gradually decreased^38^. Therefore, the decrease of the peak current density also confirmed the successful assembly of ssDNA, BSA, and target gene *hly* on the NPG/GCE electrode.

Electrochemical impedance spectroscopy (EIS) was used to further characterize the construction process of the BSA/ssDNA/NPG/GCE bioelectrode. As shown in Figure 2d, it was observed that the GCE electrode exhibited a well-defined semi-circle. After NPG loading on the GCE electrode, the diameter of the semi-circle decreased, which confirmed that the NPG exhibited excellent conductivity with improved electron transfer. This result was consistent with the higher redox peak currents of K_3_Fe(CN)_6_ for the NPG-based electrodes or bioelectrodes (Figure 2c). Subsequently, the immobilization of ssDNA on the surface of NPG provided an insulating single-strand nucleic acids layer, which obstructed electron transfer and increased resistance (Figure 2d). Further, the resistance gradually increased with the loading of BSA and the target gene *hly*. Therefore, the results of EIS provided further evidence that the BSA/ssDNA/NPG/GCE bioelectrode was successfully constructed for the detection of a nucleic acid marker (*hly* gene) and a metabolite marker (acetoin).

### Nucleic acid marker detection

Nucleic acids carry the genetic information of all known living organisms and some viruses^39^. Massive amounts of information, genetic or nongenetic, can be stored in a long piece of nucleic acids with defined sequences^39^. Some specific nucleic acid sequences carried by each species of a pathogenic microorganism can distinguish it from other organisms, and thus, the identification of bacteria or viruses can be performed by detecting the specific nucleic acid sequences^40^, as in the identification of the novel coronavirus SARS-CoV-2. Furthermore, it has been reported that the *hly* gene was used as a marker gene in *L. monocytogenes* qPCR detection^19,20^. Therefore, the *hly* gene was selected as a genetic level marker for *L. monocytogenes* detection in this study.

To confirm that the current signal was generated by MB embedded in double-stranded structure, the BSA/ssDNA/NPG/GCE bioelectrode and *hly*/BSA/ssDNA/NPG/GCE bioelectrode (incubating with 100 pM *hly* gene) were both immersed in 20 μM MB solution. After washing three times, the differential pulse voltammetry (DPV) of the BSA/ssDNA/NPG/GCE bioelectrode and *hly*/BSA/ssDNA/NPG/GCE bioelectrode were investigated in a PBS (50 mM, pH 7.0). As shown in Figure 3a, no obvious current signal was observed on the BSA/ssDNA/NPG/GCE bioelectrode. In contrast, the *hly*/BSA/ssDNA/NPG/GCE bioelectrode generated a much higher current signal. These results indicated that a large amount of MB embedded in the double-stranded structure, and the electrical signal is strong enough to detect the trace nucleic acids as shown in Figure 1b.

**Figure 3.**
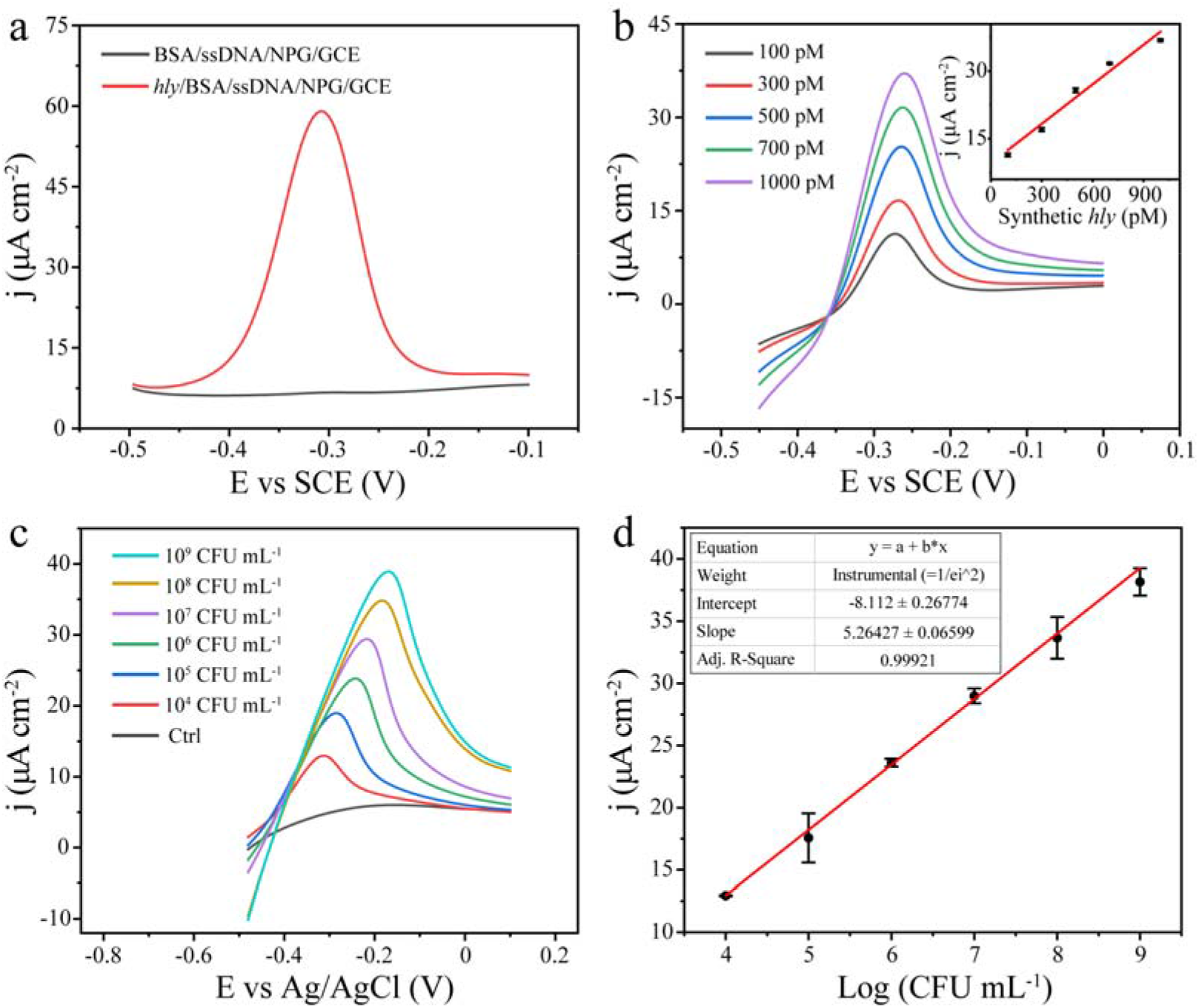
Nucleic acid marker detection. (a) The DPVs of the BSA/ssDNA/NPG/GCE electrode and *hly*/BSA/ssDNA/NPG/GCE electrode (both incubated with 20 μM MB) in a PBS (50 mM, pH 7.0). (b) The LSVs of the BSA/ssDNA/NPG/GCE bioelectrode with different concentrations of synthetic *hly* gene in PBS (50 mM, pH 7.0); inset: the linear relationship between the peak current density of MB and the synthetic *hly* gene concentrations. (c) The LSVs of the BSA/ssDNA/NPG/CWE bioelectrode with different concentrations of ultrasonically disrupted *L. monocytogenes* (10^4^~10^9^ with 10-fold increments in PBS (50 mM, pH 7.0). (d) The linear relationship between the peak current density of MB and the logarithm concentrations of *L. monocytogenes*.

To further investigate the performance of the BSA/ssDNA/NPG/GCE bioelectrode for the target gene *hly* detection, the fabricated bioelectrode was first adapted for the determination of different concentrations of synthetic *hly* gene using linear sweep voltammetry (LSV) in PBS (50 mM, pH 7.0). Figure 3b shows that the oxidation peak current of MB (at approximately −0.3 V) increased with increasing synthetic *hly* gene concentration. The observed widening and tilting of the LSV baseline signal of MB (Figure 3b) suggested that MB was undergoing redox reactions after intercalation into the double-stranded structure^41^. Moreover, the broadening electrochemical peaks were due to the gradually saturation of MB within the NPG pore volume^42^, indicating that the binding amount of MB increased with the increased formation number of double-stranded structures. The inset of Figure 3b shows that the calibration plot exhibited a good linear relationship between the peak current densities of MB and the synthetic *hly* gene concentrations in a range from 100 pM to 1000 pM. The linear regression equation was j (μA cm^−2^) = 9.52873 + 0.02926 × C_hly_ (pM) (R^2^ = 0.969) with a high sensitivity of 29.26 μA·nM^−1^·cm^−2^. In addition to the high sensitivity, the bioelecrode also had a low detection limit of 0.32 pM (S/N = 3). These results indicated that the bioelectrode had a good sensing performance for the synthetic target gene *hly* detection.

After testing the synthetic *hly* gene, the fabricated BSA/ssDNA/NPG/CWE bioelectrode on the triple functional sensing chip was used for *L. monocytogenes hly* gene detection. For this purpose, different concentrations of ultrasonically disrupted *L. monocytogenes* were used as *hly* gene samples to explore the linear relationship between the peak current densities of MB and the concentrations of *L. monocytogenes*. Figure 3c illustrates that a distinct oxidation peak of MB was clearly observed when the concentration of *L. monocytogenes* was 10^4^ CFU mL^−1^. Further, the peak current density increased with the increase in the concentration of *L. monocytogenes*. Figure 3d shows that the calibration curve had a good linear relationship between the peak current density of MB and the logarithm of *L. monocytogenes* concentration in a range from 10^4^ to 10^9^ CFU mL^−1^. The linear regression equation was expressed as follows: j (μA cm^−2^) = 5.26427 × log (CFU mL^−1^) – 8.112 (R^2^ = 0.999). The detection limit was 10^4^ CFU mL^−1^ *L. monocytogenes*. These good performances demonstrated that the rapid detection of *L. monocytogenes* was achieved using the triple functional sensing chip by detecting *hly* gene.

### Metabolite marker detection

A metabolite marker is defined as a biological characteristic that can be objectively measured and evaluated as an indicator of either normal biological processes or pathogenic processes^43^. For microorganisms, metabolite markers are intimately associated with current microbial population status, and thus, the identification of pathogenic microorganism can be achieved by detecting appropriate metabolite markers. It has been reported that acetoin can serve as a metabolite marker for the detection of *L. monocytogenes*^23^. Therefore, acetoin was selected as a metabolic level marker for *L. monocytogenes* detection in this study. The accuracy and reliability of *L. monocytogenes* detection protocols can be further improved by analyzing metabolite marker.

As a key enzyme for acetoin reduction, AR can catalyze acetoin reduce to 2,3-butanediol with NADH as a cofactor^26^, and the quantitative detection of acetoin can be achieved by measuring the consumption of NADH. Therefore, AR encoded by *bdha* gene was overexpressed and purified for acetoin detection in this study. To express AR, the *bdha* gene was amplified by PCR using *Bacillus subtilis* genomic DNA as a template. As shown in Figure S1b, an approximately 1000 bp PCR product was consistent with the *bdha* gene in size. After the construction of the recombinant pET-28a(+)-*bdha* plasmid (Figure S1a), it was transferred into *Escherichia coli* BL21(DE3). After *bdha* successful expression in *E. coli* BL21(DE3) cells, sodium dodecyl sulfate polyacrylamide gel electrophoresis (SDS-PAGE) was carried out to explore the expression level of AR (Figure S1d). Compared to *E. coli* BL21(DE3) (Figure S1d-i), the recombinant *E. coli* BL21(DE3)-AR cells produced large amounts of AR protein (Figure S1d-ii). Further, the recombinant AR protein matched the calculated molecular mass (~37 kDa) of wild-type AR from *Bacillus subtilis* strain. These results demonstrated that AR was successfully overexpressed in *E. coli* BL21(DE3) cells.

As mentioned above, the quantitative detection of acetoin could be achieved by measuring the consumption of NADH in the presence of AR. For acetoin detection, the BSA/ssDNA/NPG/CWE bioelectrode for NADH oxidation at different concentrations of acetoin was investigated using LSV in PBS(50 mM, pH 7.0) containing 3.0 mM NADH and 10 μL purified AR (5.5 mg mL^−1^). From Figure 4a, a distinct oxidation peak of NADH was obtained at a potential of approximately +0.6 V. Compared with the oxidation peak potential of NADH catalyzed using a gold sheet electrode (at approximately +0.72 V)^32^, the decrease in overpotential obviously resulted from the excellent electrocatalytic activity of NPG. As shown in Figure 4a, the peak current density of NADH decreased with the increase in acetoin concentration. The peak current density difference (Δ j) showed a strong positive correlation with the acetoin concentration in a range from 0 μM to 300 μM. The linear regression equation was Δ j (μA cm^−2^) = 0.0565 ×C_acetoin_ (μM) + 1.11558 (R^2^ = 0.998). Due to the low-coordination gold atoms contained in NPG, NPG has a good electrocatalytic activity towards NADH as mentioned above^35,36,37^. Therefore, the BSA/ssDNA/NPG/CWE bioelectrode can sensitively detect acetoin by ascertaining NADH consumption. In addition, the relationship between the quantity of *L. monocytogenes* and the concentration of acetoin was also investigated in this study. As shown in Figure 4b, the concentration of acetoin increased gradually with the increase in *L. monocytogenes* concentration. This linear correlation phenomenon was consistent with the previously reported positive correlation between acetoin concentration and *L. monocytogenes* concentration^44^. The calibration plot exhibited a strong linear relationship between the concentrations of acetoin and the logarithms of *L. monocytogenes* concentration in a range from 8.0 × 10^5^ CFU mL^−1^ to 3.5 × 10^9^ CFU mL^−1^. The linear regression equation was C_acetion_ (μM) = 87.64877 ×log (CFU mL^−1^) – 510.83867 (R^2^ = 0.970). According to these results, the triple functional sensing chip can be used to detect the concentration of acetoin by ascertaining NADH consumption with the aid of AR. The concentration of *L. monocytogenes* can then be calculated using the acetoin concentration based on the calibration plot as shown in Figure 4b.

**Figure 4.**
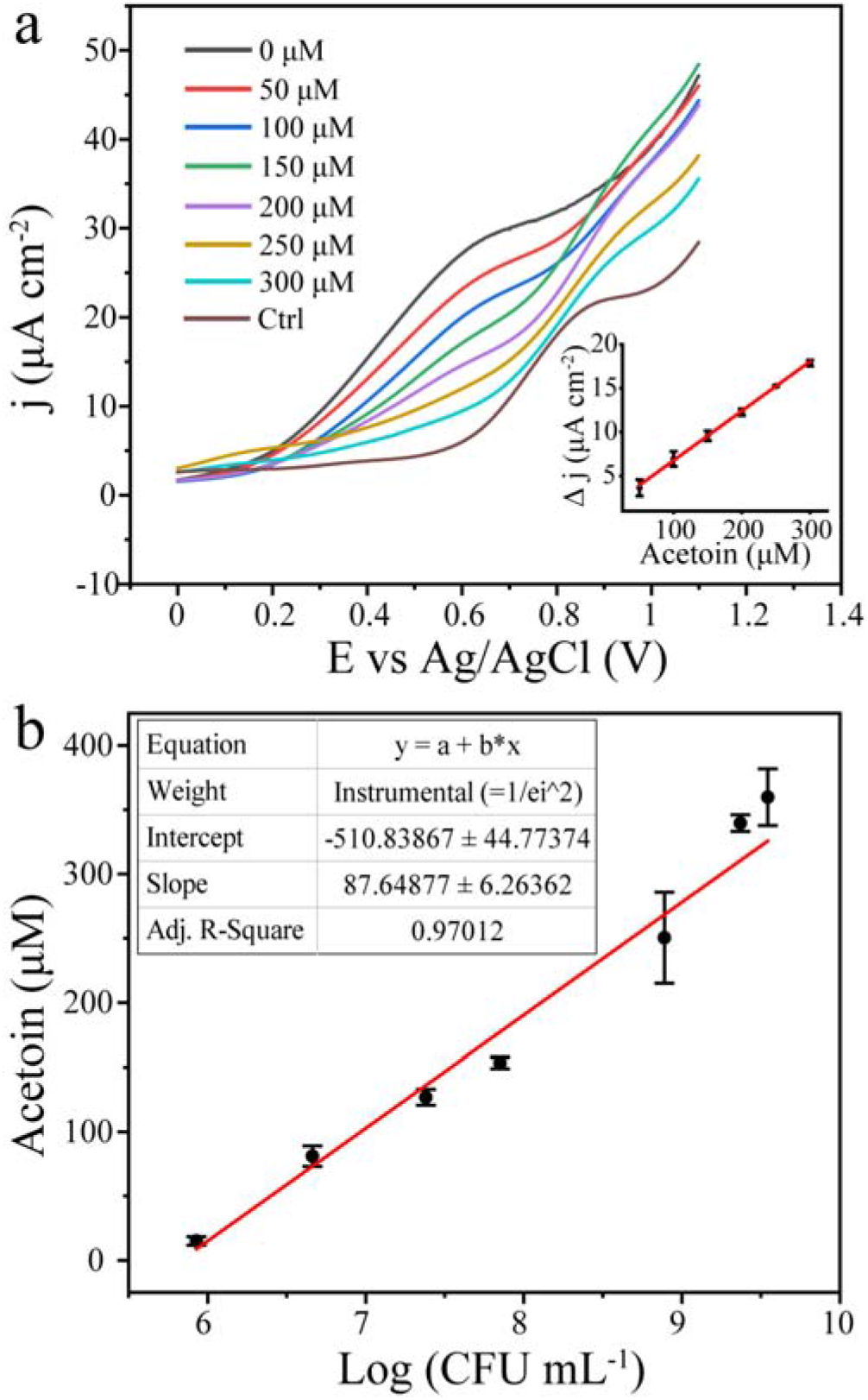
Metabolite marker detection. (a) The LSVs of the BSA/ssDNA/NPG/CWE bioelectrode in PBS (50 mM, pH 7.0) containing 3.0 mM NADH, purified AR, and different concentrations of acetoin; inset: the linear relationship between the peak current density difference (Δ j) of NADH and the acetoin concentrations. (b) The linear relationship between the concentrations of acetoin and the logarithm concentrations of *L. monocytogenes*.

### Toxin marker detection

LLO protein is a pore-forming toxin and can bind to the cholesterol of host membranes, oligomerize, and form pores to help *L. monocytogenes* escaping from the host vacuole^22^. Pore-forming phenomenon will occur when LLO protein absorbs on the membrane of Cat-Lipo. The pore-forming ability of LLO protein can be used to destruct liposome, which causes the release of liposome inclusion. If the inclusion released from liposome can be catalyzed by NPG to generate a current signal, and thus the quantitative detection of LLO protein can be realized.

To explore the pore-forming ability of the LLO protein, LLO protein was expressed and purified. As shown in Figure S2b, the *hly* gene was amplified by PCR using *L. monocytogenes* genomic DNA as a template, and an approximately 1500 bp PCR product was obtained (Figure S2b). Then, the constructed recombinant pET-28a(+)-*hly* plasmid (Figure S2a) was transferred into *E. coli* BL21(DE3). After *hly* expression, SDS-PAGE was carried out to confirm the expression level of the LLO protein (Figure S2d). Compared to *E. coli* BL21(DE3) (Figure S2d-i), the recombinant *E. coli* BL21(DE3)-LLO cells produced large amounts of LLO protein (Figure S2d-ii). Further, the recombinant LLO protein matched the calculated molecular mass (~57 kDa) of wild-type LLO protein from *L. monocytogenes* strain. These results demonstrated that LLO protein was successfully overexpressed in *E. coli* BL21(DE3) cells.

To select an ideal liposome inclusion, many substances (such as catechol, K_3_Fe(CN)_6_, hydroquinone, and etc.) were explored by the NPG/CWE electrode in a PBS (50 mM, pH 7.0) using DPV. According to Figure S3, catechol can be catalyzed by NPG, and produced a sharper oxidation peak at approximately +0.16 V with highest peak current density. Thus, catechol was selected as an ideal inclusion encapsulated into the liposome to build Cat-Lipo.

To investigate the response of the artificial Cat-Lipo to LLO protein, the Cat-Lipo was incubated with different concentrations of LLO protein, and full-wavelength scan was used to detect the absorbance of catechol released from the Cat-Lipo. Figure 5a shows that an absorbance peak at about 270 nm was clearly observed, and the absorbance value increased linearly with the increase of LLO protein concentration (Figure 5b). Besides, an obvious hole was observed on the surface of the Cat-Lipo after incubating with LLO protein (Figure 5c). These results confirmed that the pore-forming on the Cat-Lipo by LLO protein can lead to the release of catechol.

**Figure 5.**
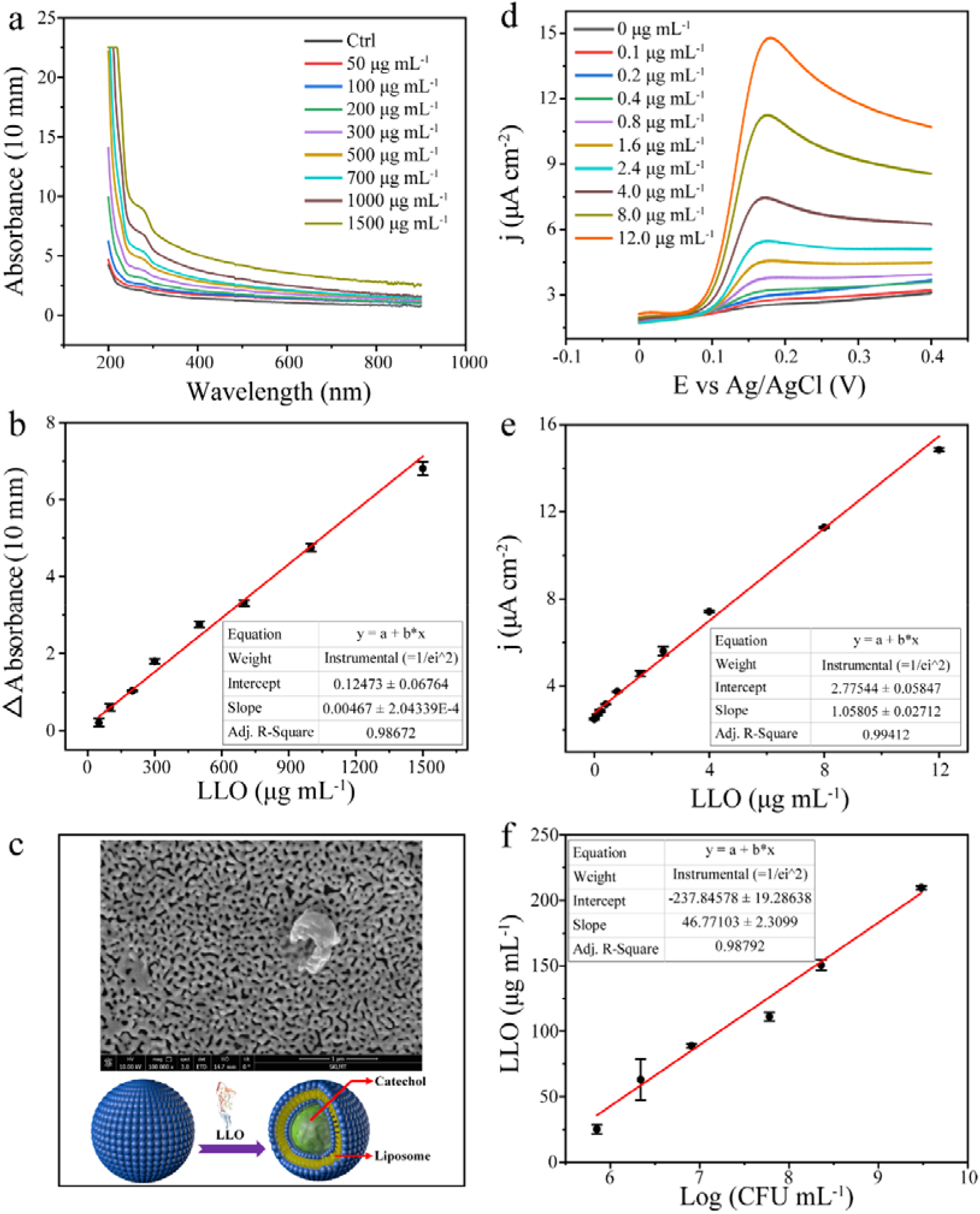
Toxin marker detection. (a) The full-wavelength scanning spectrum of the reaction mixture (Cat-Lipo incubated with different concentrations of LLO protein). (b) The linear relationship between the absorbance peak difference (Δ absorbance) and the LLO concentrations. (c) The SEM image and the three-dimensional diagram of the Cat-Lipo/NPG after incubation with the LLO protein. (d) The LSVs of the Cat-Lipo/NPG/CWE bioelectrode in a buffer (270 μL PBS (50 mM, pH 7.0) mixed with 30 μL TSB (pH 7.0)) containing different concentrations of LLO protein. (e) The linear relationship between the peak current density of catechol and the LLO concentrations. (f) The linear relationship between the concentrations of LLO and the logarithm concentrations of *L. monocytogenes*.

For the LLO protein electrochemical determination, the Cat-Lipo/NPG/CWE bioelectrodes were detected using LSV in 270 μL PBS (50 mM, pH 7.0) mixed with 30 μL Tryptic soy broth (TSB, pH 7.0) containing different concentrations of LLO protein. Figure 5d shows that the peak current density of catechol (at approximately +0.16 V) increased with the increase of LLO protein concentration. The linear regression equation was j (μA cm^−2^) = 1.05805 ×C_LLO_ (μg mL^−1^) + 2.77544 (R^2^ = 0.994) (Figure 5e). In addition, the relationship between the quantity of *L. monocytogenes* and the concentration of LLO protein was also investigated in this study. As shown in Figure 5f, the calibration plot exhibited a strong linear relationship between the concentrations of LLO protein and the logarithms of *L. monocytogenes* concentration in a range from 7.1 × 10^5^ CFU mL^−1^ to 3.0 × 10^9^ CFU mL^−1^. The linear regression equation was C_LLO_ (μg mL^−1^) = 46.77103 ×log (CFU mL^−1^) – 237.84578 (R^2^ = 0.988). According to these results, the triple functional sensing chip can be used to detect the concentration of LLO protein by detecting current produced by NPG catalysis catechol. The concentration of *L. monocytogenes* can be calculated using the LLO protein concentration based on the calibration plot as shown in Figure 5f. Therefore, the detection of *L. monocytogenes* by detecting LLO protein can be used as an auxiliary test to verify the detection results of *L. monocytogenes* via *hly* gene and acetoin detection.

### Selectivity of the triple functional sensing chip

*L. monocytogenes* was recognized as a foodborne pathogen, tolerant to extreme environmental stress conditions such as a wide temperature range (1~45°C), a broad pH range (4.5~9.6), and a high salt concentration range (10%~15%)^45^. Thus, *L. monocytogenes* may coexist with different foodborne pathogens in different types of food products. The protein, the nucleic acid sequence, and the metabolites of other pathogens may affect the selective detection of *L. monocytogenes* using the triple functional sensing chip. To explore the selectivity of the triple functional sensing chip, several microorganisms that may be present in food were used as interference bacterium such as *Bacillus cereus*^46^, *Pseudomonas aeruginosa*^47^, *Enterobacter sakazakii*^48^, *Sphingomonas paucimobilis*^49^, and *Escherichia coli*^50^. In this study, the triple functional sensing chip performed the detection of *L. monocytogenes* by integrating three different markers (*hly* or acetoin or LLO). Thus, the selectivity of the triple functional sensing chip was explored in the detection of three markers respectively.

For the selective detection of toxin marker LLO protein, the cultured *L. monocytogenes* and interference bacterium were ultrasonicated and centrifugated. Then the supernatant of *L. monocytogenes* and interference bacterium were mixed (interference bacterium: *L. monocytogenes* = 1:1) to detect using the triple functional sensing chip. From Figure 6a, the presence of the interference bacteria mentioned above caused negligible change in the relative peak current density, and the change was less than 5%. For the selective detection of nucleic acid marker *hly* gene, the mixture of bacteria (interference bacterium: *L. monocytogenes* = 1:1) was ultrasonicated, and then subjected to detect using the triple functional sensing chip. As shown in Figure 6b, it was a negligible change in the relative peak current density for the detection of *hly* gene in the presence of the interference bacteria, and the change was less than 5%. These results indicated that the triple functional sensing chip presented a strong anti-interference capability towards other interfering protein or genes because of the good blocking effect and specificity of the Cat-Lipo and capture probe. For the specificity of the triple functional sensing chip towards acetoin, a culture medium supernatant of interference bacterium was investigated via NADH consumption in the presence of AR. It is shown in Figure 6c that the interference of other bacteria was negligible in the detection of acetoin by the triple functional sensing chip. After that, the triple functional sensing chip was used to detect the culture medium supernatant of *L. monocytogenes*, and the results (Figure 6c) indicated that acetoin can be normally detected for *L. monocytogenes*. To further confirm the results, the culture medium supernatant of interference bacteria was detected by monitoring the A_340_ change of NADH in the presence of AR. As shown in Table S1, the A_340_ of NADH was almost unchanged within 10 min after adding AR. These results indicated that the triple functional sensing chip has good specificity for the detection of acetoin. Based on these results, the fabricated triple functional sensing chip can specifically detect the LLO protein, *hly* gene, and acetoin, and has potential for the detection of *L. monocytogenes* in real samples.

**Figure 6.**
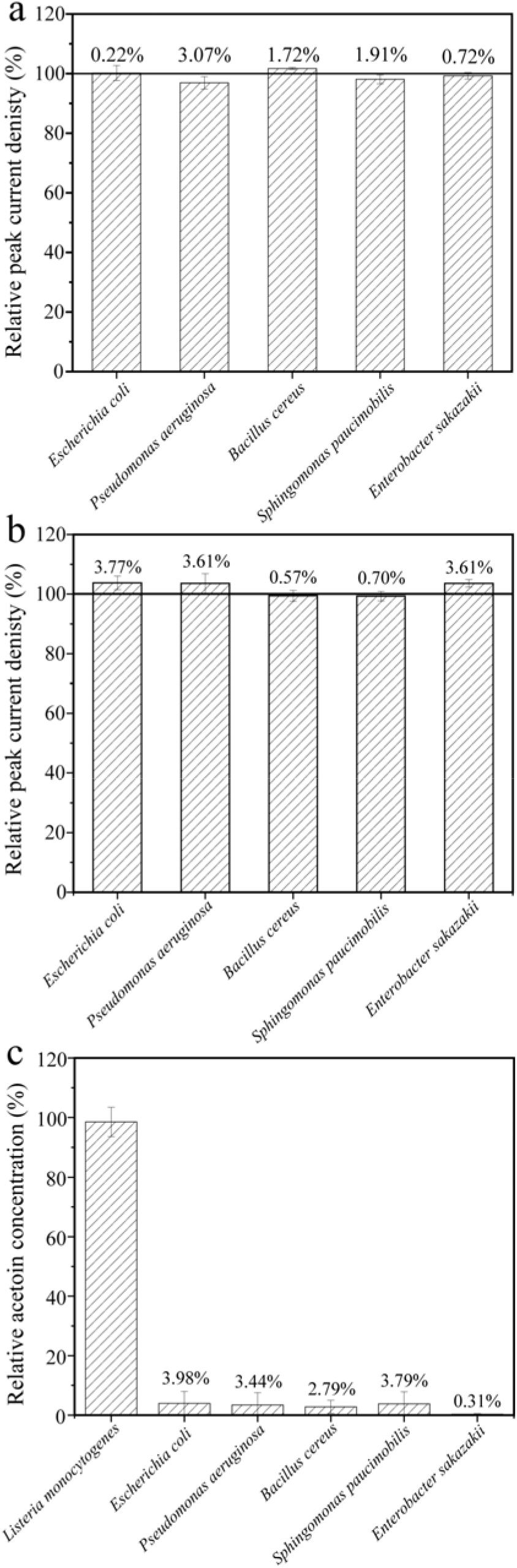
The selectivity of the triple functional sensing chip. (a) The selectivity of the triple functional sensing chip towards LLO protein (a), *hly* gene (b), and acetoin (c) (*Escherichia coli*: 3.0 × 10^9^ CFU mL^−1^; *Pseudomonas aeruginosa*: 2.0 × 10^9^ CFU mL^−1^; *Bacillus cereus*: 2.0 × 10^9^ CFU mL^−1^; *Sphingomonas paucimobilis*: 4.0× 10^8^ CFU mL^−1^; *Enterobacter sakazakii*: 2.0 × 10^9^ CFU mL^−1^).

### The detection of *L. monocytogenes* in real samples

Most countries have a zero-tolerance policy towards the presence of *L. monocytogenes* in foods due to the possible health hazards caused by this pathogen^51^. This means that pre-enrichment is required before the *L. monocytogenes* detection^52^. Further, the qualitative detection of *L. monocytogenes* is preferred in real sample analysis. To analyze the practical performance of the triple functional sensing chip, milk and instant meat were selected as samples for *L. monocytogenes* detection. The pretreatment process of the milk or instant meat sample referred to the National Food Safety Standard of China (GB 4789.30-2016). In brief, 25 mL milk or 25 g instant meat was mixed with 225 mL TSB (named solution 1), and then 0.1 mL solution 1 was added to 10 mL TSB (named solution 2). A certain amount of *L. monocytogenes* was added into the solution 2, and incubated at 37°C for 4 h, 6 h, and 8 h, respectively. After incubation, these samples were analyzed using the triple functional sensing chip and plate count method, respectively. As shown in Table 1, the concentration of *L. monocytogenes* in solution 2 can be obtained by detecting three markers (LLO or *hly* or acetoin). Compared with the plate count method (Table 1), the concentration of *L. monocytogenes* obtained by detecting LLO or *hly* or acetoin using the triple functional sensing chip had a small deviation rate (less than 8%). Based on these results, the triple functional sensing chip achieved both qualitative and quantitative detection of *L. monocytogenes*. Combined with the selectivity experimental results, the triple functional sensing chip was reliable for *L. monocytogenes* identification by simultaneously detecting a toxin marker (LLO protein), a nucleic acid marker (*hly* gene), and a metabolite marker (acetoin).

**Table 1.** The measurement of *L. monocytogenes* by the triple functional sensing chip in real samples.

| Samples | Plate count<br>(CFU mL<sup>-1</sup>) | *hly* detection (CFU<br>mL<sup>-1</sup>) | Deviation rate<br>(*hly* detection<br>vs Plate count) | Acetoin detection<br>(CFU mL<sup>-1</sup>) | Deviation rate<br>(Acetoin<br>detection vs<br>Plate count) | LLO detection<br>(CFU mL<sup>-1</sup>) | Deviation rate<br>(LLO<br>detection vs<br>Plate count) |
| --- | --- | --- | --- | --- | --- | --- | --- |
| Milk | $9.40 \times 10^7$ | $(9.89 \pm 1.37) \times 10^7$ | +5.21 % | $(9.26 \pm 3.07) \times 10^7$ | -1.49 % | $(9.05 \pm 0.02) \times 10^7$ | -3.72 % |
| | $7.50 \times 10^8$ | $(7.71 \pm 0.97) \times 10^8$ | +2.80 % | $(7.49 \pm 1.47) \times 10^8$ | -0.13 % | $(7.37 \pm 0.16) \times 10^8$ | -1.73 % |
| | $2.10 \times 10^9$ | $(2.06 \pm 0.08) \times 10^9$ | -1.90 % | $(2.19 \pm 0.91) \times 10^9$ | +4.29 % | $(2.25 \pm 0.19) \times 10^9$ | +7.14 % |
| Instant meat | $7.30 \times 10^7$ | $(7.62 \pm 0.72) \times 10^7$ | +4.38 % | $(7.16 \pm 2.16) \times 10^7$ | -1.92 % | $(7.68 \pm 0.95) \times 10^7$ | +5.21 % |
| | $1.40 \times 10^8$ | $(1.33 \pm 0.11) \times 10^8$ | -5.00 % | $(1.41 \pm 2.82) \times 10^8$ | +0.71 % | $(1.36 \pm 0.07) \times 10^8$ | -2.86 % |
| | $8.90 \times 10^8$ | $(9.13 \pm 0.26) \times 10^8$ | +2.58 % | $(9.10 \pm 0.35) \times 10^8$ | +2.24 % | $(8.76 \pm 0.26) \times 10^8$ | -1.57 % |

## Discussion

In summary, the rapid and accurate detection of *L. monocytogenes* was achieved using a triple functional sensing chip by simultaneously detecting a toxin marker (LLO protein), a nucleic acid marker (*hly* gene), and a metabolite marker (acetoin). Compared with the conventional culture-based gold standard method, the proposed triple functional sensing chip does not require technical expertise and complex identification procedures such as dynamic test, catalase test, sugar fermentation, and hemolytic properties. Meanwhile, the long identification procedure time of 2~5 days was reduced to about 90 min using the triple functional sensing chip. Further, the results obtained from the detection of *L. monocytogenes* at different levels (LLO at the protein level, *hly* at the genetic level, and acetoin at the metabolic level) using the triple functional sensing chip can be mutually verified, which reduces the probability of false positives based on a single marker (protein or nucleic acid or metabolite). As for pathogenic microorganisms (such as the novel coronavirus SARS-CoV-2), detection that only relies on a single marker, requires repeating detection multiple times to avoid a false positives result. The integrated sensing chip constructed in this study may provide a novel solution for the detection of pathogenic microorganisms.

## Methods

### Materials and reagents

Acetoin, phosphatidylcholine, cholesterol, and 1-octadecanethiol were purchased from Shanghai Macklin Biochemical Co., Ltd. (China). MB was obtained from Tianjin Bodi Chemical Co., Ltd. (China). NADH-Na_2_, imidazole, and GeneGreen (a nucleic acid fluorescent dye) were purchased from Beijing Dingguo changsheng Biotechnology Co., Ltd. (China). Isopropyl-b-D-thiogalactoside (IPTG), kanamycin sulfate (Kana), sodium dodecyl sulfate (SDS), and BSA were purchased from Pengyuan Biotechnology Co., Ltd. (China). PBS (50 mM) was prepared using Na_2_HPO_4_·12H_2_O and NaH_2_PO_4_·2H_2_O. All other reagents used were of analytical grade. Ultrapure water (> 18 MΩ cm) was used throughout all the experiments. Lysis equilibration buffer (LE buffer, pH 8.0) was prepared with 7.8 g L^−1^ NaH_2_PO_4_·2H_2_O and 17.54 g L^−1^ NaCl. Luria-Bertani (LB, pH 7.0) was prepared with 10.0 g L^−1^ tryptone, 5.0 g L^−1^ yeast extract, and 10.0 g L^−1^ NaCl. TSB (pH 7.0) was composed of 17.0 g L^−1^ tryptone, 3.0 g L^−1^ soy peptone, 2.5 g L^−1^ glucose, 5.0 g L^−1^ NaCl, and 2.5 g L^−1^ K_2_HPO_4_. The thiolated capture probe was synthesized by Suzhou Genewiz Biotechnology Co., Ltd. (China) (sequence: 5’-TGGCGGCACATTTGTCACTGCA-(CH2)_3_-SH-3’). The 48-base *hly* sequence originated from *L. monocytogenes hly* gene was synthesized by Sangon Biotech Co., Ltd. (China) (sequence: 5’-TGCAGTGACAAATGTGCCGCCAAGAAAAGGTTATAAAGATGGAAATGA-3’).

### Bacterial strains, plasmids, and culture conditions

*L. monocytogenes* was cultivated in TSB at 37°C in a gyratory shaker at 200 rpm. *E. coli* BL21(DE3) was used for heterologous expression of AR or LLO and routinely grown in LB medium at 37°C. *Bacillus subtilis* was incubated in LB at 37°C in a gyratory shaker at 200 rpm. All strains used in this study were preserved in 17% (v/v) glycerin solution, and then stored at −80°C. pET-28a(+) (Beijing Dingguo Changsheng Biotechnology Co., Ltd., China) was used as the expression vector for AR or LLO heterologous expression in *E. coli* BL21(DE3).

### Heterologous expression of LLO and AR

For LLO expression, *hly* gene was amplified by PCR using *L. monocytogenes* genomic DNA as a template. The following primers were used: 5’-CGC<u>GGATCC</u>ATGAAAAAAATAATGCTAGTTTTTATTACAC-3’ (forward, BamHI site underlined) and 5’-CCG<u>CTCGAG</u>TTCGATTGGATTATCTACACT-3’ (reverse, XhoI site underlined). The procedure of heterologous expression referred to our previous report^31^. For AR expression, the *bdha* gene was amplified by PCR using *Bacillus subtilis* genomic DNA as a template. The following primers were used: 5’-CGC<u>GGATCC</u>ATGAAGGCAGCAAGATGGCATAAC-3’ (forward, BamHI site underlined) and 5’-CCG<u>CTCGAG</u>GTTAGGTCTAACAAGGATTTTGACTTGG-3’ (reverse, XhoI site underlined). The procedure of heterologous expression referred to our previous report^31^. After expression, the *E. coli* BL21(DE3) cells expressed LLO or AR were collected by centrifugation at 6000 rpm for 10 min, and washed with PBS (50 mM, pH 7.0). The pellet was then resuspended in LE buffer (pH 8.0), and broken by high pressure homogenizer (SPCH-18, Stansted Fluid Power Products Ltd., UK). Additionally, to examine the *hly* or *bdha* gene expression in *E. coli* BL21(DE3) cells, the collected cells were lysed with pyrolysis and electrophoretically using 12% (w/v) SDS-PAGE. After that, LLO or AR purified by Ni-NTA (GenScript Biotech Co., China) procedure was eluted using imidazole buffers (pH 8.0) with different concentrations. According to previous report^53^, the catalytic activity of AR was measured by monitoring the substrate-dependent absorbance change of NADH at 340 nm (A_340_) using NanoPhotometer (N60, Implen Inc., Germany). One unit (U) of enzyme activity was defined as the amount of enzyme required to oxidize 1 μmol of NADH per minute.

### Preparation of Cat-Lipo/NPG/CWE bioelectrode

NPG was fabricated according to our previous report^11^. A paper sensor with four CWE, one CCE, and one reference electrode (Ag/AgCl) was used as the multi-biocatalyst sensing chip. The CWE was coated by NPG to build the NPG/CWE electrode. The electrochemical active areas of these NPG/CWE electrodes were detected by CV in a 0.5 M H_2_SO_4_ aqueous solution at a scan rate of 50 mV s^−1^.

The Cat-Lipo used for electrode modification were prepared by a probe sonicator. Phosphatidylcholine, cholesterol, and 1-octadecanethiol were mixed in chloroform in a small vial (molar ratio 10:10:1). The organic solvent was then removed with a N_2_ stream to form a thin lipid layer on the bottom. Appropriate amount of 0.5 M catechol solution was added so that the final lipid concentration was 1.0 mg mL^−1^, followed by sonicating for 10 min in a bath sonicator. Then the solution was sonicated for 1 min at 60 W by a probe sonicator (JY92-IIDN, Ningbo Scientz Biotechnology Co., Ltd., China). After that, the Cat-Lipo was collected by centrifugation, and washed with PBS (50 mM, pH 7.0) three times to remove free catechol. The pellet was then resuspended in PBS (50 mM, pH 7.0) to a final concentration of 0.08 mg μL^−1^. Then 40 μL Cat-Lipo (0.08 mg μL^−1^) was dropped onto the surface of NPG/CWE electrode to build the Cat-Lipo/NPG/CWE bioelectrode.

### Preparation of BSA/ssDNA/NPG/CWE bioelectrode

The prepared NPG/CWE electrodes were immersed in an ssDNA solution (1 μM) at 4°C for 72 h. The resulting ssDNA/NPG/CWE bioelectrodes were washed with PBS (50 mM, pH 7.0) three times to remove unconjugated capture probes. The ssDNA/NPG/CWE bioelectrodes were then immersed in 1% (w/v) BSA at room temperature for 30 min to block the nonspecific binding sites, and the prepared BSA/ssDNA/NPG/CWE bioelectrodes were washed with PBS (50 mM, pH 7.0) three times and stored at 4°C for use.

### Experimental measurements

The morphology of Cat-Lipo/NPG was characterized using a SEM (Nova NanoSEM 450, FEI Co. USA). After loading ssDNA and *hly* gene, the *hly*/ssDNA/NPG composite and NPG were incubated with GeneGreen (a nucleic acid fluorescent dye), washed three times, and then imaged at 470 nm blue light wavelength. To facilitate the study of electrode interface characteristics, some experiments (bioelectrode CV characteristics, EIS analysis, MB signal generating by MB embedded in double-stranded structure, and synthetic *hly* detection) were performed in a three-electrode cell at room temperature using a CHI 760E electrochemical workstation (Shanghai Chenhua Apparatus Co., Ltd. China). A glassy carbon electrode (GCE) modified by NPG was used as a working electrode for further modification, and a Pt sheet (1 cm × 1 cm) and a saturated calomel electrode (SCE) were used as counter electrode and reference electrode, respectively. Besides, all other electrochemical detections (*L. monocytogenes hly* gene detection, acetoin detection, LLO detection, anti-interference, and real samples analysis) were performed on the triple functional sensing chip at room temperature using a CHI 760E electrochemical workstation (Shanghai Chenhua Apparatus Co., Ltd. China). As an electrolyte, PBS (50 mM) was deoxygenated by N_2_ for 20 min before used. All experiments were repeated at least three times.

### Standard plate count method

The standard plate count method was used to verify the amount of *L. monocytogenes*. After serial dilution, samples were plated on solid medium (TSB with 18 g L^−1^ agar added) and incubated at 37°C for 24 h to count colonies. The colonies of *L. monocytogenes* on TSB plates were recorded.

### The detection of LLO protein

The culture of *L. monocytogenes* incubated in TSB overnight at 37°C at 200 rpm was diluted to 10^5^ CFU mL^−1^ as an inoculum. The medium inoculated with diluted culture was incubated in a gyratory shaker at 37°C at 200 rpm. At regular time intervals, samples were removed from the medium to determine the colony numbers of *L. monocytogenes* and LLO concentration. The colony numbers of these samples were analyzed using the standard plate count method. Additionally, these samples were subjected to sonication for 15 min using a sonicator Q125 (Qsonica Co., USA) and centrifugated to obtained the supernatant. The LLO in supernatant was detected by monitoring the A_270_ change of catechol released from the Cat-Lipo. The reaction mixtures containing 90 μL supernatant and 10 μL Cat-Lipo (0.08 mg μL^−1^). The reaction occurred at room temperature for 4 h. The A_270_ change values (ΔA_270_) were substituted into the prepared calibration curve (Δ A_270_ vs LLO concentration) to calculate the concentration of LLO. To further investigated the LLO concentration, the supernatant was analyzed by 12% SDS-PAGE. The band gray value at 57 kDa was compared with the gray value of standard BSA band to calculated the LLO concentration.

For LLO protein electrochemical determination, the Cat-Lipo/NPG/CWE electrode was used as a working electrode. 270 μL PBS (50 mM, pH 7.0) mixed with 30 μL TSB (pH 7.0) containing different concentrations of LLO were added onto the sensing chip, and incubated for 90 min at room temperature. The peak currents were detected using LSV (20 mV s^−1^) technique.

### The detection of *hly* gene

For *hly* gene detection, the cultured *L. monocytogenes* was washed with PBS and diluted to different concentrations (10^4^~10^9^ CFU mL^−1^). Then 1.0 mL of *L. monocytogenes* samples with different concentrations (10^4^~10^9^ CFU mL^−1^) were denatured by heating in a metal bath (95°C) for 10 min and immediately chilled in ice to obtain denatured single-strand DNA^54^. These samples were subjected to sonication for 15 min using a sonicator Q125 (Qsonica Co., USA) to break the long DNA strands into smaller fragments. The BSA/ssDNA/NPG/CWE bioelectrode was incubated with the DNA fragments for 10 min at room temperature to construct a *hly*/BSA/ssDNA/NPG/CWE bioelectrode. The *hly*/BSA/ssDNA/NPG/CWE bioelectrode was immersed in a 20 μM MB solution at room temperature for 5 min. In order to remove free MB, the obtained bioelectrodes were then washed with 0.5% (w/v) SDS solution and PBS (50 mM, pH 7.0) for three times, respectively. Then LSV (20 mV s^−1^) was used to detect the oxidation peak current of MB embedded in double-stranded structure. A linear relationship between the oxidation peak current densities of MB and the logarithms of *L. monocytogenes* concentration was constructed.

### Acetoin detection

The culture of *L. monocytogenes* incubated in TSB overnight at 37°C at 200 rpm was diluted to 10^5^ CFU mL^−1^ as an inoculum. The medium inoculated with diluted culture was incubated in a gyratory shaker at 37°C at 200 rpm. At regular time intervals, samples were removed from the medium to determine the colony numbers of *L. monocytogenes* and acetoin concentration. The colony numbers of these samples were analyzed using the standard plate count method. Additionally, the supernatant was collected after centrifugation at 5000 rpm for 10 min, and the concentration of acetoin in the supernatant was detected by monitoring the A_340_ change of NADH. The reaction mixtures containing 94 μL supernatant, 3 μL of 100 mM NADH, and 3 μL of 5.5 mg mL^−1^ AR. The reaction occurred at room temperature for 10 min. The A_340_ change values (Δ A_340_) were substituted into the prepared calibration curve (Δ A_340_ vs acetoin concentration) to calculate the concentration of acetoin. For acetoin electrochemical detection, the BSA/ssDNA/NPG/CWE bioelectrode was used to detect the consumption of NADH using LSV in PBS (50 mM, pH 7.0) containing 3.0 mM NADH. Different concentrations of acetoin and 10 μL of 5.5 mg mL^−1^ AR were added into PBS buffer, and then the mixture reacted at room temperature for 10 min. PBS (50 mM, pH 7.0) containing 3.0 mM NADH served as a control. After reaction, the BSA/ssDNA/NPG/CWE bioelectrode was used to detect the decrease in the oxidation peak current of NADH. A linear relationship between the differences in the oxidation peak current density of NADH and the concentrations of acetoin was constructed.

### Anti-interference of the triple functional sensing chip

In this study *Bacillus cereus*, *Escherichia coli*, *Pseudomonas aeruginosa*, *Sphingomonas paucimobilis*, and *Enterobacter sakazakii* were used as interference bacterium. These bacteria were cultured in LB at 37°C in a gyratory shaker. The concentrations of these bacteria were obtained using the standard plate count method.

For the anti-interference analysis of LLO or *hly* detection, the interference bacteria were mixed with *L. monocytogenes* (bacterium concentration ratio = 1:1). The LLO or *hly* detection was performed as the mentioned above. For the anti-interference analysis of acetoin detection, synthetic sample (60 μL culture medium supernatant of the interference bacterium and 240 μL PBS (50 mM, pH 7.0)) containing 3.0 mM NADH was used as a buffer. The acetoin detection procedure was the same as mentioned above.

## Acknowledgments

This work was supported by grants from National Key Research and Development Program of China (2019YFA0904800), National Natural Science Foundation of China (32070097 and 91951202), and Shandong Science & Technology Fund Planning Project (2019GSF109069).

## Author contributions

Yachao Zhang: Investigation, writing-original draft. Huimin Wang: Visualization. Sa Xiao: Formal analysis. Xia Wang: Conceptualization, supervision, resources, writing-review & editing. Ping Xu: Resources, discussion& writing-review & editing.

## References

1. Murray, E. G. D., Webb, R. A., & Swann, M. B. R. A disease of rabbits characterised by a large mononuclear leucocytosis, caused by a hitherto undescribed bacillus *bacterium monocytogenes*. J. Pathol. Bacteriol. 29, 407–439 (1926).

2. Schlech, W. F. et al. Epidemic Listeriosis-evidence for transmission by food. New Engl. J. Med. 308, 203–206 (1983).

3. Hamon, M., Bierne, H. & Cossart, P. *Listeria monocytogenes*: a multifaceted model. Nat. Rev. Microbiol. 4, 423–434 (2006).

4. Radoshevich, L. & Cossart, P. *Listeria monocytogenes*: towards a complete picture of its physiology and pathogenesis. Nat. Rev. Microbiol. 16, 32–46 (2018).

5. Soni, D. K., Ahmad, R. & Dubey, S. K. Biosensor for the detection of *Listeria monocytogenes*: emerging trends. Crit. Rev. Microbiol. 44, 590–608 (2018).

6. Banada, P. P. et al. Optical forward-scattering for detection of *Listeria monocytogenes* and other *Listeria* species. Biosens. Bioelectron. 22, 1664–1671 (2007).

7. Etty, M. C. et al. New immobilization method of anti-PepD monoclonal antibodies for the detection of *Listeria monocytogenes* p60 protein - Part B: Rapid and specific sandwich ELISA using antibodies immobilized on a chitosan/CNC film support. React. Funct. Polym. 143, 104317 (2019).

8. Ikeda, M., Yamaguchi, N. & Nasu, M. Rapid on-chip flow cytometric detection of *Listeria monocytogenes* in milk. J. Health Sci. 55, 851–856 (2009).

9. Datta, A. R., Moore, M. A., Wentz, B. A. & Lane, J. Identification and enumeration of *Listeria monocytogenes* by nonradioactive DNA probe colony hybridization. Appl. Environ. Microbiol. 59, 144–149 (1993).

10. Paul, M., Baranzoni, G. M., Albonetti, S. & Brewster, J. D. Direct, quantitative detection of *Listeria monocytogenes* in fresh raw whole milk by qPCR. Int. Dairy J. 41, 46–49 (2015).

11. Zhang, Y. et al. Electrochemical immunosensor for HBe antigen detection based on a signal amplification strategy: The co-catalysis of horseradish peroxidase and nanoporous gold. Sens. Actuators B: Chem. 284, 296–304 (2019).

12. Robinson, A. L., Lee, H. J. & Ryu, D. Polyvinylpolypyrrolidone reduces cross-reactions between antibodies and phenolic compounds in an enzyme-linked immunosorbent assay for the detection of ochratoxin A. Food Chem. 214, 47–52 (2017).

13. Resendiz-Nava, C., Esquivel-Hernandez, Y., Alcaraz-Gonzalez, A., Castaneda-Serrano, P. & Nava, G. M. PCR assays based on *invA* gene amplification are not reliable for *Salmonella* detection. Jundishapur J. Microbiol. 12, e68764 (2019).

14. Attar, A. et al. Amperometric inhibition biosensors based on horseradish peroxidase and gold sononanoparticles immobilized onto different electrodes for cyanide measurements. Bioelectrochemistry 101, 84–91 (2015).

15. Cho, S., Yang, H. C. & Rhee, W. J. Simultaneous multiplexed detection of exosomal microRNAs and surface proteins for prostate cancer diagnosis. Biosens. Bioelectron. 146, 111749 (2019).

16. Zhao, K. et al. Simultaneous detection of three biomarkers related to acute myocardial infarction based on immunosensing biochip. Biosens. Bioelectron. 126, 767–772 (2019).

17. Drummond, T. G., Hill, M. G. & Barton, J. K. Electrochemical DNA sensors. Nat. Biotechnol. 21, 1192–1199 (2003).

18. Niu, X. et al. Electrochemical DNA biosensor based on gold nanoparticles and partially reduced graphene oxide modified electrode for the detection of *Listeria monocytogenes hly* gene sequence. J. Electroanal. Chem. 806, 116–122 (2017).

19. Volokhov, D., Rasooly, A., Chumakov, K. & Chizhikov, V. Identification of *Listeria* Species by microarray-based assay. J. Clin. Microbiol. 40, 4720–4728 (2002).

20. Rodriguez-Lazaro, D. et al. Quantitative detection of *Listeria monocytogenes* and *Listeria innocua* by real-time PCR: Assessment of *hly*, *iap*, and *lin02483* targets and amplifluor technology. Appl. Environ. Microbiol. 70, 1366–1377 (2004).

21. Hamon, M. A., Ribet, D., Stavru, F. & Cossart, P. Listeriolysin O: the swiss army knife of *Listeria*. Trends Microbiol. 20, 360–368 (2012).

22. Morton, C. J., Sani, M., Parker, M. W. & Separovic, F. Cholesterol-dependent cytolysins: membrane and protein structural requirements for pore formation. Chem. Rev. 119, 7721–7736 (2019).

23. Yu, Y. X., Sun, X. H., Liu, Y., Pan, Y. J. & Zhao, Y. Odor fingerprinting of *Listeria monocytogenes* recognized by SPME-GC-MS and E-nose. Can. J. Microbiol. 61, 367–372 (2015).

24. Zhu, Y. et al. Mesoporous tungsten oxides with crystalline framework for highly sensitive and selective detection of foodborne pathogens. J. Am. Chem. Soc. 139, 10365–10373 (2017).

25. Zhu, Z. et al. Cr doped WO3 nanofibers enriched with surface oxygen vacancies for highly sensitive detection of the 3-hydroxy-2-butanone biomarker. J. Mater. Chem. A 6, 21419–21427 (2018).

26. Molinnus, D. et al. Development and characterization of a field-effect biosensor for the detection of acetoin. Biosens. Bioelectron. 115, 1–6 (2018).

27. Romeo, A., Leung, T. S. & Sánchez, S. Smart biosensors for multiplexed and fully integrated point-of-care diagnostics. Lab Chip 16, 1957–1961 (2016).

28. Sage, A. T., Besant, J. D., Lam, B., Sargent, E. H. & Kelley, S. O. Ultrasensitive electrochemical biomolecular detection using nanostructured microelectrodes. Accounts Chem. Res. 47, 2417–2425 (2014).

29. Wang, X. et al. Enzyme-nanoporous gold biocomposite: Excellent biocatalyst with improved biocatalytic performance and stability. PLoS One 6, e24207 (2011).

30. Wu, C., Liu, Z., Sun, H., Wang, X. & Xu, P. Selective determination of phenols and aromatic amines based on horseradish peroxidase-nanoporous gold co-catalytic strategy. Biosens. Bioelectron. 79, 843–849 (2016).

31. Liu, Z. et al. Highly sensitive microbial biosensor based on recombinant *Escherichia coli* overexpressing catechol 2,3-dioxygenase for reliable detection of catechol. Biosens. Bioelectron. 126, 51–58 (2019).

32. Qiu, H. et al. Enzyme-modified nanoporous gold-based electrochemical biosensors. Biosens. Bioelectron. 24, 3014–3018 (2009).

33. Rohs, R., Sklenar, H., Lavery, R. & Röder, B. Methylene blue binding to DNA with alternating GC base sequence: A modeling study. J. Am. Chem. Soc. 122, 2860–2866 (2000).

34. Nicholson, W. L. The *Bacillus subtilis ydjL* (*bdhA*) gene encodes acetoin reductase/2,3-butanediol dehydrogenase. Appl. Environ. Microbiol. 74, 6832–6838 (2008).

35. Fujita, T., Qian, L. H., Inoke, K., Erlebacher, J. & Chen, M. W. Three-dimensional morphology of nanoporous gold. Appl. Phys. Lett. 92, 251902 (2008).

36. Fujita, T. et al. Atomic origins of the high catalytic activity of nanoporous gold. Nat. Mater. 11, 775–780 (2012).

37. Molina, L. M. & Hammer, B. Theoretical study of CO oxidation on Au nanoparticles supported by MgO(100). Phys. Rev. B 69, 155424 (2004).

38. Li, Y. et al. A dual-type responsive electrochemical immunosensor for quantitative detection of PCSK9 based on n-C_60_-PdPt/N-GNRs and Pt-poly (methylene blue) nanocomposites. Biosens. Bioelectron. 101, 7–13 (2018).

39. Ke, Y., Castro, C. & Choi, J. H. Structural DNA nanotechnology: Artificial nanostructures for biomedical research. Annu. Rev. Biomed. Eng. 20, 375–401 (2018).

40. Li, Y. et al. Detection and identification of genetic material via single-molecule conductance. Nat. Nanotechnol. 13, 1167–1173 (2018).

41. Veselinovic, J., Almashtoub, S. & Seker, E. Anomalous trends in nucleic acid-based electrochemical biosensors with nanoporous gold electrodes. Anal. Chem. 91, 11923–11931 (2019).

42. Kelley, S. O., Barton, J. K., Jackson, N. M. & Hill, M. G. Electrochemistry of methylene blue bound to a DNA-modified electrode. Bioconjugate Chem. 8, 31–37 (1997).

43. Naylor, S. Biomarkers: Current perspectives and future prospects. Expert Rev. Mol. Diagn. 3, 525–529 (2003).

44. Romick, T. L. & Fleming, H. P. Acetoin production as an indicator of growth and metabolic inhibition of *Listeria monocytogenes*. J. Appl. Microbiol. 84, 18–24 (1998).

45. Vallim, D. C. et al. Twenty years of *Listeria* in Brazil: Occurrence of *Listeria* species and *Listeria monocytogenes* serovars in food samples in Brazil between 1990 and 2012. Biomed Res. Int. 2015, (2015).

46. Forghani, F. Wei, S. & OH, D. H. A rapid multiplex real-time PCR high-resolution melt curve assay for the simultaneous detection of *Bacillus cereus*, *Listeria monocytogenes*, and *Staphylococcus aureus* in food. J. Food Protect. 79, 810–815 (2016).

47. Werner, B. G., Wu, J. Y. & Goddard, J. M. Antimicrobial and antifouling polymeric coating mitigates persistence of *Pseudomonas aeruginosa* biofilm. Biofouling 35, 785–795 (2019).

48. Friedemann, M. *Enterobacter sakazakii* in food and beverages. Int. J. Food Microbiol. 116, 1–10 (2007).

49. Adekoya, I. et al. Occurrence of bacteria and endotoxins in fermented foods and beverages from Nigeria and South Africa. Int. J. Food Microbiol. 305, 108251 (2019).

50. Scott, M. E. et al. *Salmonella* and shiga toxin-producing *Escherichia coli* in products sampled in the food safety and inspection service raw pork baseline study. J. Food Protect. 83, 552–559 (2020).

51. Upadhyay, A. et al. Inactivation of *Listeria monocytogenes* on frankfurters by plant-derived antimicrobials alone or in combination with hydrogen peroxide. Int. J. Food Microbiol. 163, 114–118 (2013).

52. Cambero, M. I. et al. Sanitation of selected ready-to-eat intermediate-moisture foods of animal origin by e-beam irradiation. Foodborne Pathog. Dis. 9, 594–599 (2012).

53. Fu, J. et al. Metabolic engineering of *Bacillus subtilis* for chiral pure meso-2,3-butanediol production. Biotechnol. Biofuels 9, 90 (2016).

54. Pandey, C. M., Singh, R., Sumana, G., Pandey, M. K. & Malhotra, B. D. Electrochemical genosensor based on modified octadecanethiol self-assembled monolayer for *Escherichia coli* detection. Sens. Actuators B: Chem. 151, 333–340 (2011).

