## Supplementary information for "A triple functional sensing chip for rapid detection of pathogenic *Listeria monocytogenes*"

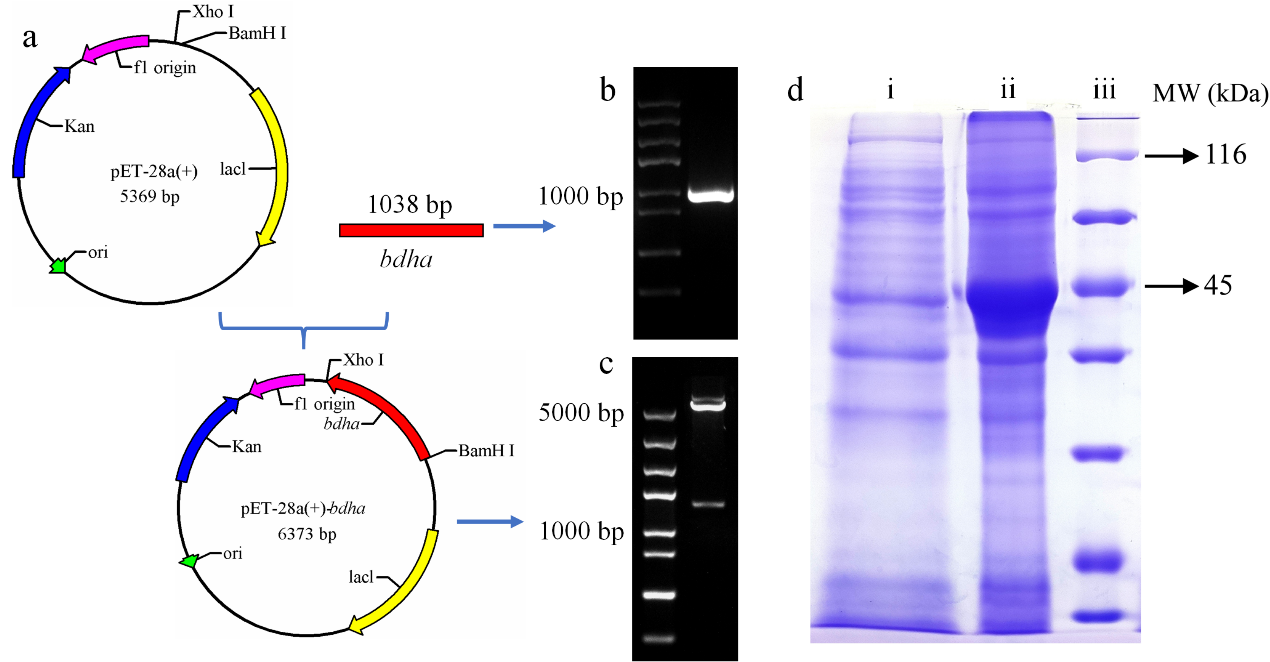

**Figure S1** The recombinant expression of acetoin reductase in *E. coli* BL21(DE3). (a) The schematic illustration of a recombinant expression vector construction. (b) The *bdha* gene PCR product. (c) The expression vector of pET-28a(+)-*bdha* digested with BamHI and XhoI. (d) The SDS-PAGE of *E. coli* BL21(DE3)-AR (lane ii) and *E. coli* BL21(DE3) (lane i); line iii was protein molecular weight marker.

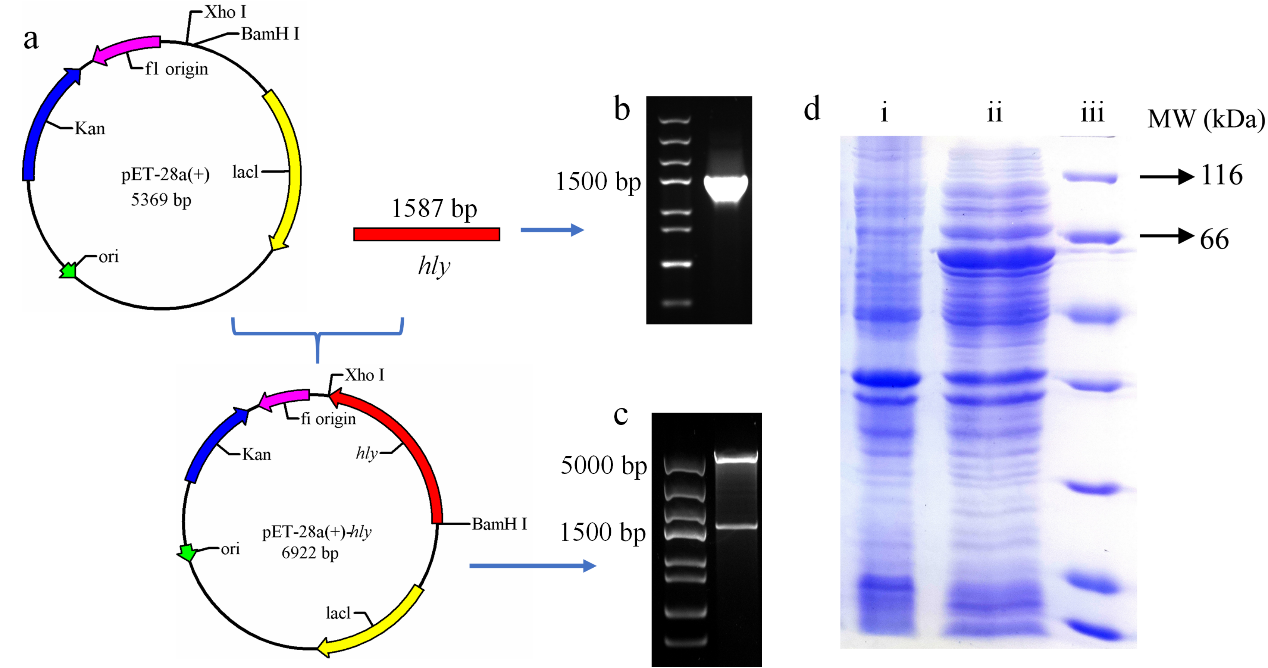

**Figure S2** The recombinant expression of LLO in *E. coli* BL21(DE3). (a) The schematic illustration of a recombinant expression vector construction. (b) The *hly* gene PCR product. (c) The expression vector of pET-28a(+)-*hly* digested with BamHI and XhoI. (d) The SDS-PAGE of *E. coli* BL21(DE3)-LLO (lane ii) and *E. coli* BL21(DE3) (lane i); line iii was protein molecular weight marker.

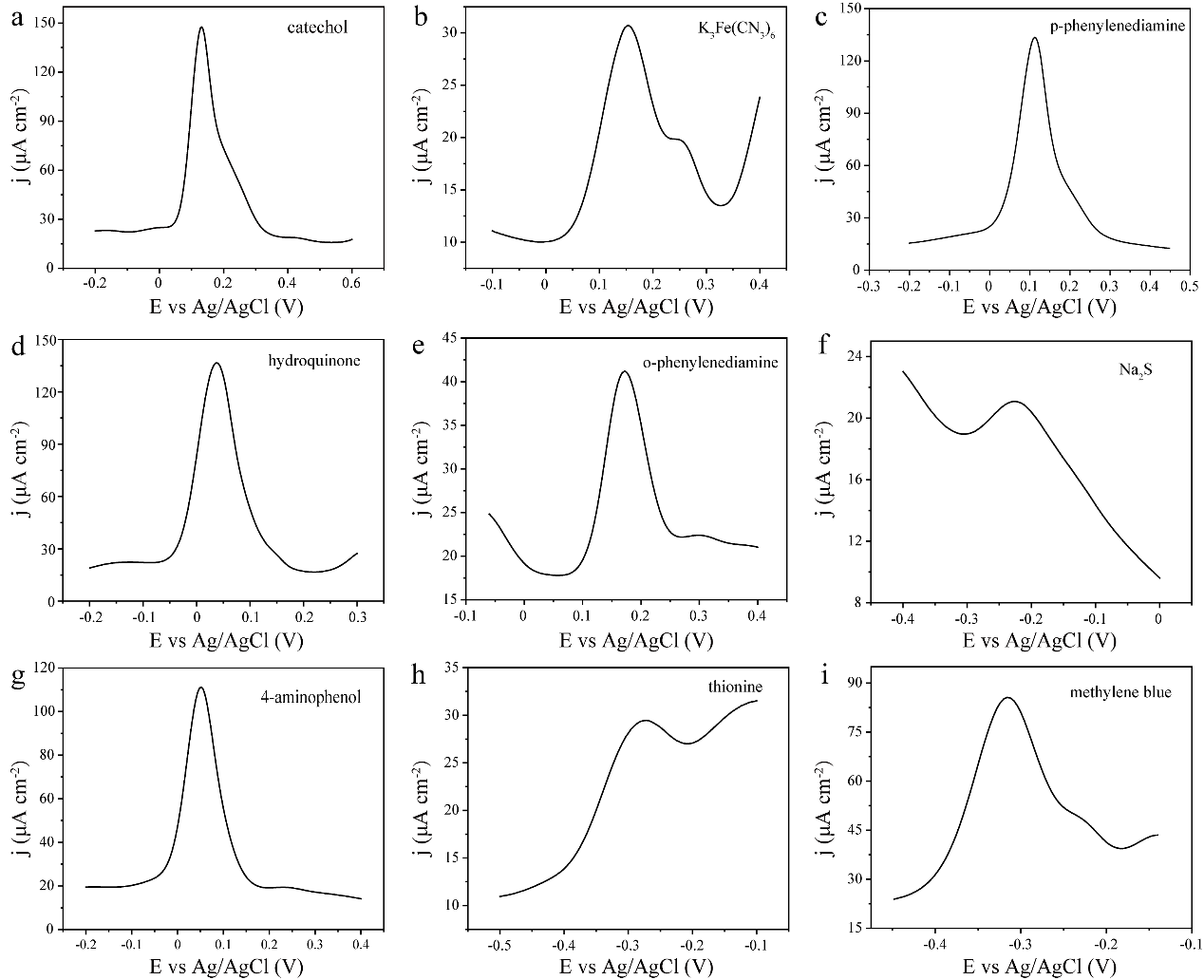

**Figure S3** The DPVs of the NPG/CWE electrode in a PBS (50 mM, pH 7.0) containing different substances (200 μM).

**Table S1** The A_340_ of NADH after adding AR in bacterium supernatant.

| Bacterium strains | Adding AR 0 min (A_340_) | Adding AR 10 min (A_340_) |
| --- | --- | --- |
| *Escherichia coli* | 18.88 | 18.83 |
| *Pseudomonas aeruginosa* | 18.87 | 18.73 |
| *Bacillus cereus* | 18.47 | 18.42 |
| *Sphingomonas paucimobilis* | 19.48 | 19.37 |
| *Enterobacter sakazakii* | 21.70 | 21.48 |
